# Deliberation is a controllable process governed by desirability and cognitive effort

**DOI:** 10.64898/2025.12.12.693733

**Authors:** Jan A. Calalo, Seth R. Sullivan, Nicholas R. Muscara, John Buggeln, Truc T. Ngo, Matthew R. Short, Michael J. Carter, Isaac L. Kurtzer, Joshua G.A. Cashaback

## Abstract

Humans spend a lifetime making decisions based on incoming sensory information and goals. Prominent theories of perceptual decision-making have described the components of deliber-ation process, yet they lack a unifying principle that governs how the nervous system tunes the deliberation process across multiple contexts. Desirability (reward) and effort (energy) are major determinants in governing a broad range of human and animal behaviour, such as for-aging, walking, and decisions. Here we develop a theory where desirability and cognitive effort tune the control gains that govern deliberation. Several hallmark features of decision behaviour simply emerge from the model, with the deliberation process closely resembling low-dimensional neural dynamics. We also predict and provide a novel mechanistic explanation for choking-under-pressure, where extremely large rewards lead to performance deficits. Our principled framework explains both behavioural and neural phenomena while providing a path to unify disparate fields.

## Introduction

> *He moves through space with minimum waste and maximum joy*
>
> — Sade

Decision-making is ubiquitous in our everyday lives, from the cognitively effortful choices of an air traffic controller to selecting between delicious food options. Prominent perceptual decision-making theories have typically viewed decision deliberation through an information processing perspective,^1^ such as drift-diffusion models, that act as input-output systems (**Fig. 1A, light grey only**). These theories have significantly advanced our understanding of deliberation and replicated several decision-making phenomena.^1,2^ Yet they lack a unifying principle that governs *how* the nervous system sets the parameters (gains) for different experimental contexts, such as reward changes or speed instructions. Intuitively, we prioritize sensory information linked to desirable and rewarding outcomes, and have diminished focus on sensory information when tasks are effortful and offer little reward. Building on past work, here we propose a novel theoretical framework that deliberation is a controllable process governed by desirability and cognitive effort (**Fig. 1A, dark plus light grey**). By considering desirability and cognitive effort as the underlying objectives that govern deliberation, several hallmark features of behaviour and neural phenomena simply emerge from our model across multiple contexts.

**Figure 1:**
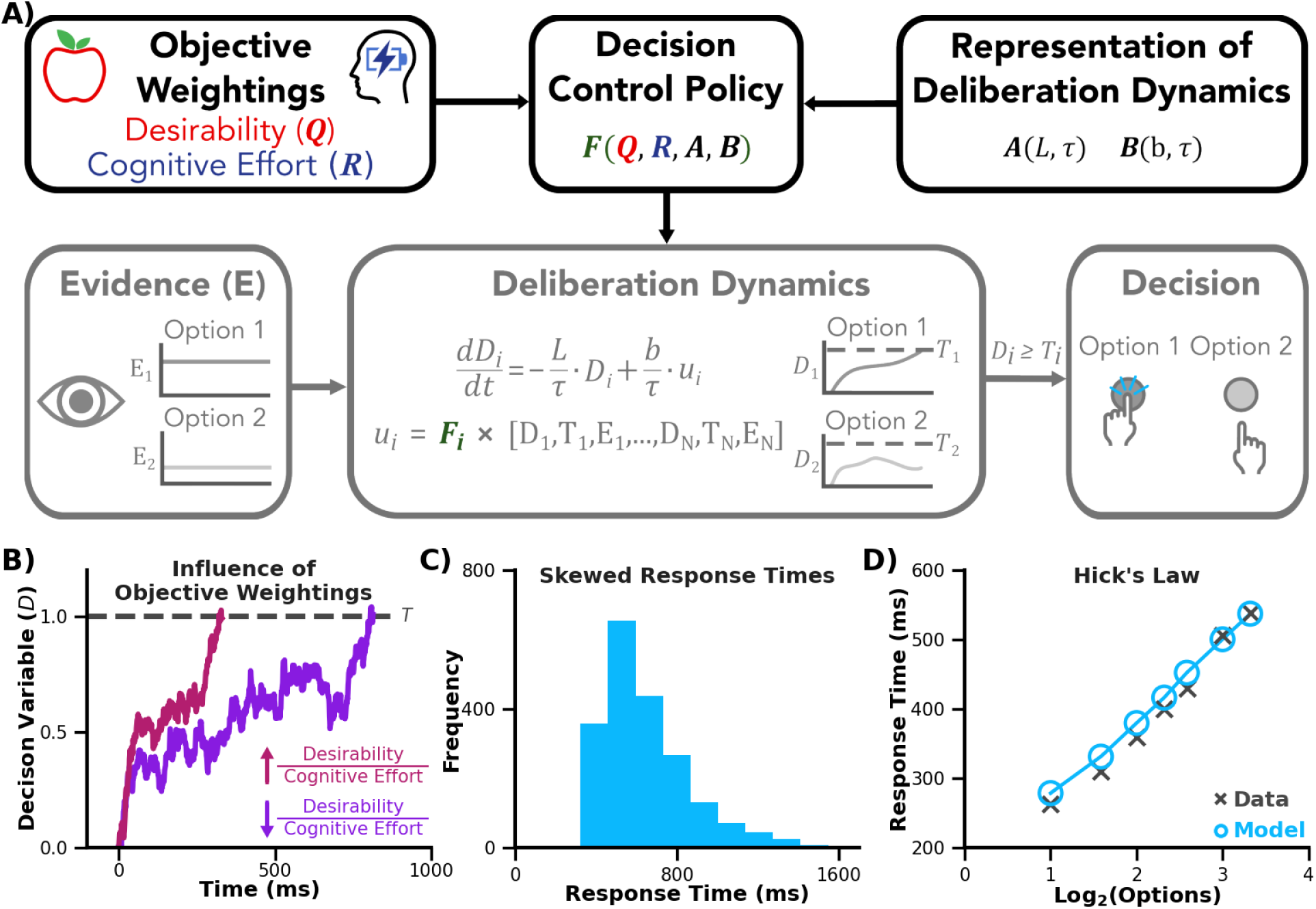
Desirability and cognitive effort dictate decision-making. **A)** Our novel theory (dark gray + light gray) builds upon an information processing perspective (light gray only) by viewing deliberation as a controllable process. We propose that the **objectives** of desirability (*Q*) and cognitive effort (*R*), along with a **representation of deliberation dynamics** (e.g., some knowledge of leak: *L*), influences a **decision control policy**. **Evidence** (*E*) feeds into the **deliberation dynamics**. The evolution of the decision variable (*D*) is determined by control gains (*F*) set by the decision control policy. The idea that a control policy determines the control gains on deliberation dynamics is in stark contrast to the information processing perspective, which does not explain *how* the gains (i.e., free parameters) are set by the nervous system. The control gains govern the neural control signal (*u*) used to drive the decision variable for a particular option (*i*). A **decision** is made once a decision variable is greater than or equal to a decision threshold (*T*). **B)** The influence of desirability and cognitive effort on deliberation. Given the same level of evidence, a high desirability to cognitive effort ratio (reddish purple) leads to faster decisions, while comparatively a low desirability to cognitive effort ratio (purple) leads to slower decisions. From our theory several hallmark features of decision behaviour emerge, including **(C)** skewed response times, **(D)** Hick’s law [adapted from Hick 1952], and more (**Fig. 2 & 4**), as well as explaining low-dimensional neural data in a new light (**Fig. 3 & 4**).

## Results

### Deliberation as a Controllable Process

It is well-established that the nervous system cares about the objectives of maximizing desirability^3^ and minimizing cognitive effort.^4,5^ Indeed, reward and punishment influence the desirability of different options and consequently the deliberation.^3,6^ Here we consider cognitive effort to be the neural control signal that drives the deliberation process. Cognitive effort, which is energetically costly,^7^ influences several cognitive processes including decision-making.^4^ More generally, energy is a major determinant of behavior, from cognition^8^ to walking^9^ to foraging.^10^ Together, desirability and cognitive effort are biologically plausible objectives that have been shown to influence a host of natural behaviours.

Extensive research has shown that the nervous system forms representations to predict and generate future behaviour. For example, it is well-established that the sensorimotor system uses a representation of body dynamics to make predictions of upcoming consequences of neural activity and muscle forces.^11,12^ For decision-making to utilize objectives like desirability and cognitive effort, the nervous system would have to have some representation of deliberation dynamics.

We propose that the objectives of desirability and cognitive effort, as well as a representation of deliberation dynamics, determine a decision control policy with control gains. To formalize this concept, we use optimal control theory^13,14^ (linear-quadratic regulator; see **Methods**). Here the idea is that the nervous system sets control gains that maximize desirability and minimize cognitive effort during decision-making. Critically, the control gains determine the strength of the neural control signal that drives the decision variable.

The input to deliberation is sensory information, notably goal-related evidence. A decision variable reflects the ongoing deliberation, and is both moved by a neural control signal and integrated over time. A neural control signal is a function of the control gains, which determine i) how evidence is accumulated, and ii) the urgency to make a decision. To accumulate evidence, momentary evidence is multiplied by a control gain. This is similar to standard drift-diffusion models where some “gain” parameter, often termed drift rate, is multiplied by evidence and integrated over time.

We also consider the urgency to make a decision. That is, an evidence-independent process is used to make a decision, a notion supported by behaviour^2^ and neural recordings.^15,16^ The difference between the current decision variable state and the decision threshold is used to drive the decision variable. This process acts to push a decision variable towards a threshold, resulting in a growing and evidence-independent component of deliberation. While our usage of urgency differs from some accounts where an urgency signal is multiplied by low-pass filtered evidence,^2,17^ it aligns with others suggesting that urgency is an additive process.^15,16^

Information processing models, such as drift-diffusion, urgency, or combinations thereof, act as input-output systems where parameters are fit to the experimental data. For example, in the drift-diffusion model the gain parameter that multiplies evidence is fit to the data. Critically, this class of models does not explain *how* these parameters are set. The novelty of our approach is that the control gains are tuned based on biologically plausible objectives and a representation of deliberation dynamics. This enables our model to accumulate more evidence and have greater urgency when there is high desirability and low cognitive effort (**Fig. 1B, reddish purple**). Conversely, the model will accumulate less evidence and have lower urgency when there is low desirability and high cognitive effort (**Fig. 1B, purple**). That is, novelly and unlike past models, the objectives of the nervous system modulate the relative roles of evidence accumulation and urgency.

There are a few remaining important details of the model. We use a parallel racer formulation to simultaneously consider different options.^18^ We also consider leak (*L*),^18^ a temporal constant (*τ*),^2,17^ and state noise (*σ*).^2,17,18,19,20,21,22^ Lastly, *b* transforms the neural control signal to move the decision variable.

Decisions are typically expressed behaviourly with some motor action, such as a button push, a saccade, or a reach to a target. In our model, a decision is made once a decision variable is greater than or equal to a decision threshold. The corresponding output of the model is response time and option selection.

Our model predicts several hallmark features of perceptual decision-making (**Fig. 1C,D; Fig 2**). By accounting for noise in the deliberation process, we predict skewed response times (**Fig. 1C**). We also replicate Hick’s law, where an increase in the number of options leads to a logarithmic increase in response times (**Fig. 1D**).^23^ Hick’s law arises through lateral inhibition, which is the increase of the decision variable of one option leading to a decrease in the decision variable of the other options.^18^ We find lateral inhibition emerges from the control policy by considering the states of the parallel options while optimizing the cost composed of desirability and cognitive effort.

**Figure 2:**
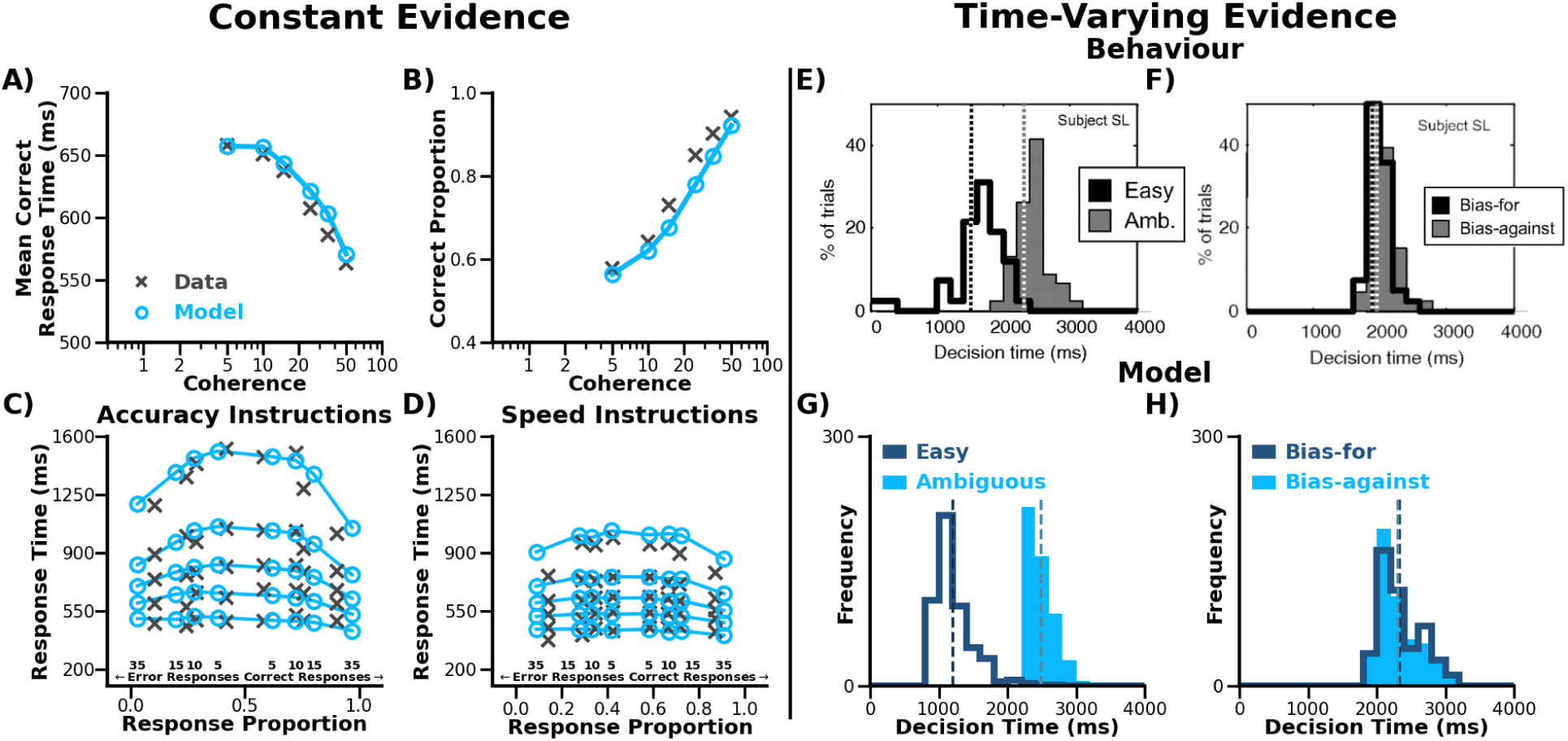
Replication of Decision-Making Phenomena A-D) The control policy acts on evidence as a function of the desirability and cognitive effort. Participant behaviour (grey x’s) from the random-dot motion discrimination task showing **A)** a decrease in mean response time of correct responses and **B)** an increase in correct responses with stronger evidence (i.e., higher coherence) [Adapted from Ratcliff, 2008]. Our model (light blue circles) captures these trends by selecting control gains that consider the objectives of desirability and cognitive effort. **C-D)** When instructed to make fast decisions or accurate decisions, humans have been shown to change their decision-making behaviour (grey x’s) [Adapted from Ratcliff, 2008]. Our model (light blue circles) replicates behavioural changes between different task contexts due to task instructions by modulating the desirability of making a decision. We model the speed instructions using a higher desirability of making a decision. The higher desirability results in more error decisions and faster decision times. We model the accuracy instructions using a lower desirability resulting in more correct but relatively slower decisions. **E-H)** Urgency emerges from the control policy and is needed to explain behaviour with time-varying evidence. Participants observed 15 tokens move individually into one of two targets. They were instructed to select the target that would finish with the most tokens. **E-F)** Histograms of decision times in the tokens task [Adapted from Cisek, 2009]. **E)** With the easy token patterns, the tokens had a high probability of moving into one of the targets compared to the ambiguous token patterns. Humans made faster decisions with easy token patterns. **F)** In the bias-for and bias-against token patterns, the first 3 tokens respectively went into the same or the opposite target as the final correct option. Participants did not have different decision times in the bias-for and bias-against decisions. In our model that considers urgency, we were also able to replicate the **(G)** different decision times between easy and ambiguous token patterns and the **(H)** similar decision times between bias-for and bias-against token patterns. Collectively, our model can capture stereotypical decision behaviour related to constant and time-varying evidence.

### Acting on Evidence Based on Desirability and Cognitive Effort

Next, we show that our novel theory can capture several important perceptual decision-making phenomena.^19,24,25,26^ In the random dot motion discrimination task, participants are asked to determine the overall motion direction of a cloud of moving dots. The percentage of dots that move in the same direction is experimentally manipulated and termed coherence. To model the resulting decision-making behaviour, prior computational theories fit model parameters to each level of evidence.^19,21^ In contrast, we use constant desirability, cognitive effort, and other parameters to determine a single set of control gains for any level of evidence. Our model uses these control gains to predict faster (**Fig. 2A**) and more accurate decisions (**Fig. 2B**) with higher coherence, aligning with past literature.^19^

It is widely known that the speed-accuracy tradeoff is influenced by task context (i.e., experimental conditions). When participants are instructed to make fast decisions, they make faster but less accurate decisions compared to when asked to make accurate decisions (**Fig. 2C, D**).^19^ It is natural to think that it is more desirable to select an option more quickly with the fast instructions. But since prior theories do not consider high-level objectives, they fit different gain parameters between conditions to replicate the speed-accuracy tradeoff. Conversely, our model can capture these different task contexts by modulating desirability that result in changes of the control gains. Higher desirability leads to a stronger urge to make a decision resulting in faster and more incorrect decisions (**Fig. 2D**). Thus, a key feature of our work is that we provide an explicit relationship between the changes in objectives and the mechanisms that drive deliberation.

### Urgency Arises from Desirability and Cognitive Effort

The random dot coherence task typically uses constant evidence where the coherence is fixed within a trial. Cisek and colleagues (2009) developed the tokens task that used time-varying evidence that changes within a trial. Participants observed tokens move individually into one of two targets and were instructed to select the target they felt would finish with the most tokens. This task uses time-varying evidence since the correct target probability changes over time. Participants decided faster with easy token patterns when tokens moved with a high probability to one of the targets compared to ambiguous token patterns when tokens moved more randomly to either target (**Fig. 2E**). For bias-for and bias-against token patterns, the initial evidence went respectively towards or away from the correct target. Participants had the same decision times for the bias-for and bias-against token patterns (**Fig. 2F**). Evidence accumulation alone would find earlier decisions in the bias-for token patterns compared to the bias-against token patterns. Hence, the authors proposed that urgency was critical in explaining the behavioural results.

As a reminder, in our model urgency is the difference between the deliberation variable state and the decision threshold, multiplied by a control gain. We can replicate the decision times found with dynamic evidence (**Fig. 2G, H**) because urgency emerges from the control policy that trades off the objectives of desirability and cognitive effort.

### Neural Activity

Our theory can also capture neural activity observed during decision-making tasks. Steinemann and colleagues (2024) measured neural activity in the lateral intraparietal sulcus of non-human primates during the random dot motion discrimination task.^16^ Using dimensionality reduction techniques, they found low-dimensional neural activity related to the deliberation of a particular target (**Fig. 3A**). They proposed that the neural activity could be decomposed into evidence-independent (urgency) (**Fig. 3A**, grey line only) and evidence-dependent (evidence accumulation) components (**Fig. 3B**). The evidence accumulation component is found by subtracting out the urgency component from the neural activity. In our model, the decision variable aligns closely with the low-dimensional neural activity (**Fig. 3D**).^16,25^ We replicate the evidence accumulation component by subtracting out the contribution of urgency (**Fig. 3E**).

**Figure 3:**
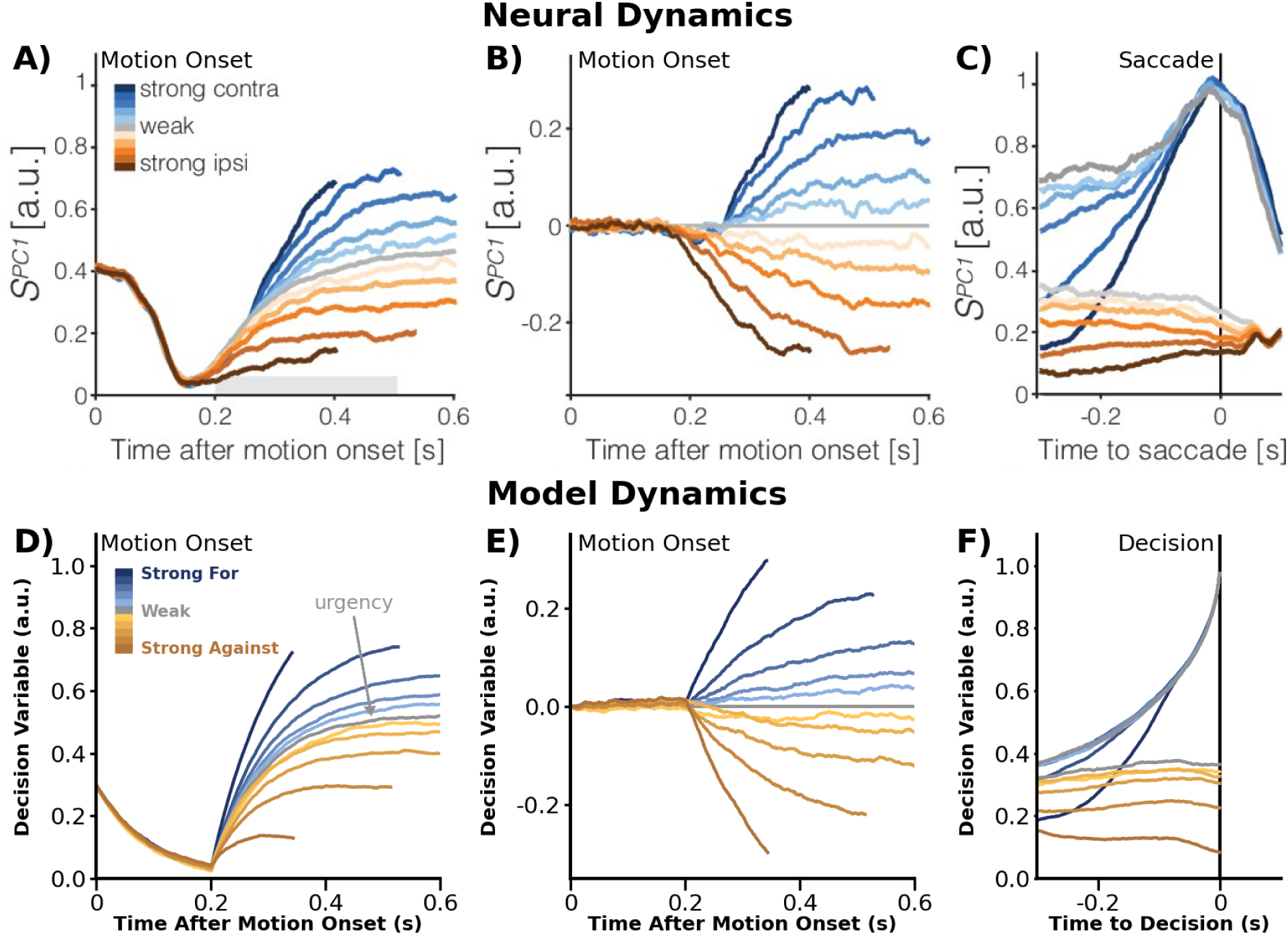
Neural activity has evidence-dependent and evidence-independent components. A-C) Low-dimensional neural activity in the lateral intraparietal sulcus of primates during a random dot motion discrimination task [Adapted from Steinemann, 2024]. Grey represents 0% coherence (weak evidence). Various shades of blue and orange represent different coherence levels towards and away from the tuning direction of the neural population, respectively. **A)** Average neural activity aligned to motion onset. **B)** Evidence-dependent component of the neural activity for the corresponding data shown in **A**, found by subtracting the average activity during the 0% coherence condition (grey). **C)** Average neural activity temporally aligned to a saccade. **D-F)** Simulated model dynamics with different levels of coherence. **D)** Average simulated decision variable over time of one potential option for different coherence levels. **E)** Evidence-dependent component of the simulated decision variable for the corresponding decision variable shown in **D**, found by subtracting the average simulated decision variable during the 0% coherence condition (grey). **F)** Average simulated decision variable temporally aligned with the time of decision. The decision variable of our model does well to capture temporally evolving low-dimensional neural activity.

When the researchers temporally aligned the neural activity to the saccade indicating a decision, the neural activity overlaps as it crossed some threshold (**Fig. 3C**). When we temporally align our model simulations to the decision time, we can see similar trends where the decision variable converges as it approaches the decision (**Fig. 3F**). Compatible with the neural data, the model makes decisions with no evidence by crossing some fixed decision threshold due to urgency, rather than some collapsing decision threshold as used by many drift-diffusion models.^20,27,28^

### Choking-Under-Pressure

A critical feature of our model is that the desirability of a decision influences the deliberation. Recent work with human and non-human primates shows that reward and performance follows an inverted-U pattern, where extremely large (jackpot) rewards leads to “choking-under-pressure” (**Fig. 4A**).^29^ Corresponding low-dimensional neural activity in non-human primates was projected onto target location and reward axes (**Fig. 4B**), where neural activity farther from the center corresponded to successfully hitting the target (**Fig. 4B pink panel**). In line with prospect theory,^30^ the value of a decision is relative to some subjective reference value. Specifically, our model can capture this choking-under-pressure phenomenon by modulating the relative desirability of a potential option compared to the average expected value (**Fig. 4C**). Medium and high rewards cause a more ideal balance of urgency and evidence accumulation leading to more success. Conversely, a jackpot reward, where the relative reward is very high, leads to a greater reliance on sensory evidence accumulation but a detrimental reduction in urgency. Here the idea is that one wants to reduce urgency to both accumulate more evidence and avoid inaccurate decisions. However, decreasing urgency too much results in a lower decision variable and thus less decisions under a time constraint. The lowered decision variable in the jackpot condition aligns with the lowered neural activity found in the experimental data (**Fig. 4B,D**). We suggest that the mechanism leading to the neural bias found by the researchers is due to a tradeoff between evidence accumulation and urgency as a function of relative desirability. Thus, our theory helps bridge perceptual decision-making with neuroeconomics and sensorimotor control.

**Figure 4:**
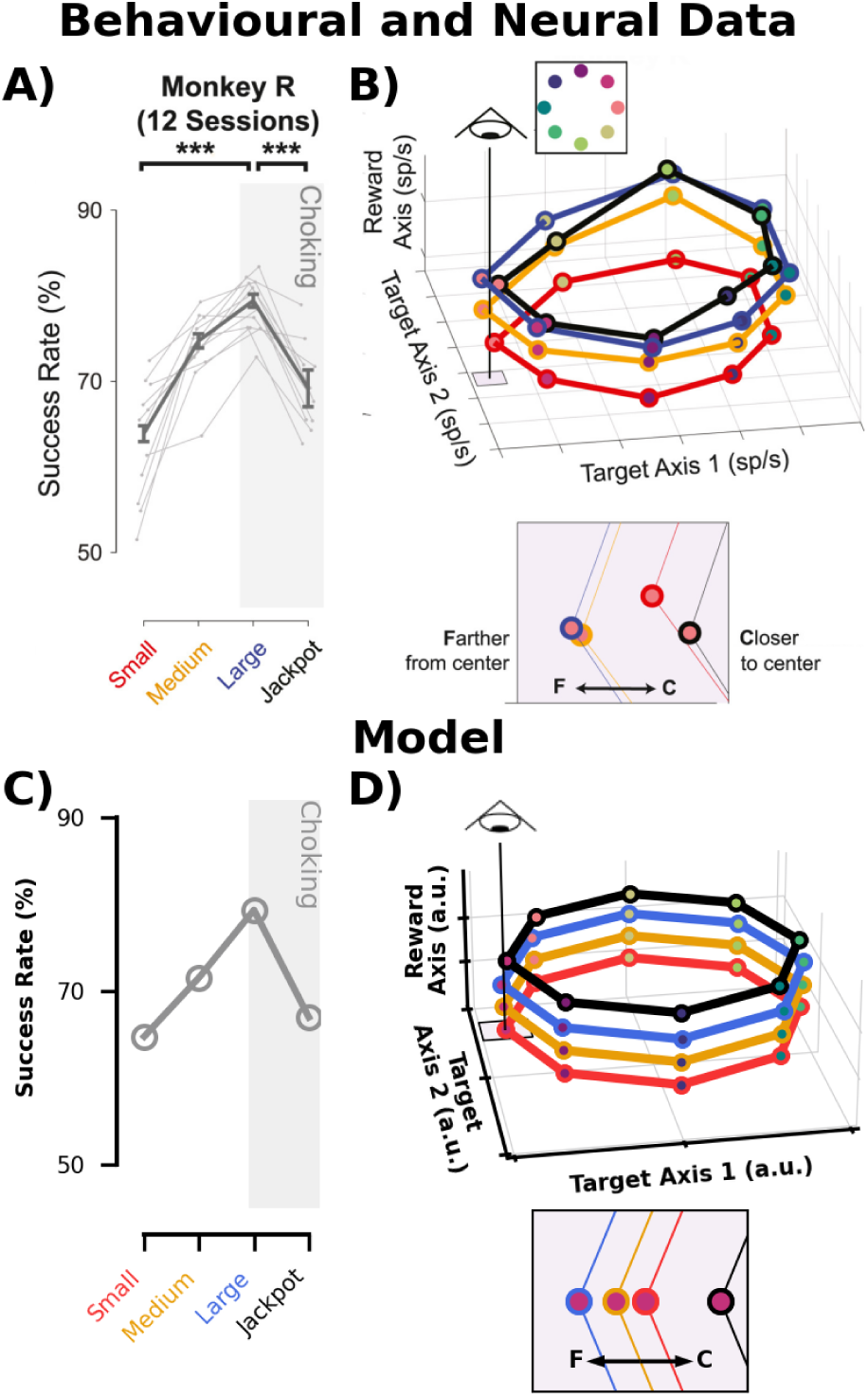
Relative desirability ‘explains choking-under-pressure’ phenomena. Smoulder and colleagues (2024) measured the neural activity related to “choking-under-pressure” in monkeys when reaching to multiple potential targets.^29^ Immediately prior to each reach, the monkeys were informed which target to select and the corresponding juice reward (small, medium, large, “jackpot”) if they hit the target. After a very brief delay that allowed some deliberation, the monkeys were cued to initiate their reach. **A-B)** Experimental and neural data [Adapted from Smoulder, 2024]. The colors represent the level of reward for the trial. **A)** Success rate given different reward magnitudes. Here, success rate was defined as the percent of reaches that hit a target. The decrement in performance in the jackpot condition compared to the large reward condition demonstrates the ‘choking-under-pressure’ phenomenon. **B)** Neural activity at the time of the movement cue, which is projected projected onto reward and target location axes. The pink panel shows a zoomed in view of the neural projections for each level of reward, relative to the center of the target location plane. Neural activity farther from center was associated with more successful reaches. **C-D)** Simulated behavioural and neural data. **C)** Success rate given different reward magnitudes. Here, success rate is the percentage of trials where the correct target was selected by the end of the trial. **D)** The average decision variable across each option projected onto the target location and level of reward. By considering desirability within a trial as a function of relative reward, we replicate that a jackpot results in worse performance.

## Discussion

Our novel theory builds upon prior work by proposing that the gains which control deliberation are determined by the objectives of desirability and cognitive effort. With our computational framework, we can capture hallmark behavioural and neural phenomena across experimental contexts.

In our framework, the control policy for deliberation is determined by the higher level objectives of desirability and cognitive effort. There is ample evidence across foraging, motor, and cognitive behaviours that humans and animals consider a tradeoff between reward and movement related metabolic energy.^9,10^ In neuroeconomics, cognitive effort relates to metabolic energy,^31^ where blood glucose levels decrease during cognitively demanding tasks.^32^ Here we propose that there is a tradeoff with reward and cognitive effort during perceptual decision-making. Future work should manipulate and measure cognitive effort to better understand its influence on perceptual decision-making.

We found that the decision variable over time closely resembled low-dimensional neural activity. Steinemann and colleagues (2024) found the neural activity in the lateral intraparietal region corresponds to ongoing deliberation.^16^ They found signals related to evidence accumulation and urgency, which emerge from our computational decision-making framework. The basal ganglia have been implicated with both reward and urgency.^33^ This is consistent with our framework where urgency arises from a cost function which includes desirability. Premotor and primary motor cortices have been shown to correspond with ongoing deliberation between multiple potential reaching targets,^34^ which is compatible with the idea that deliberation influences neural activity in the motor cortex to explain choking-under-pressure.^29^ It would be interesting to examine the influence of desirability and cognitive effort on the embodiment of deciding and acting.

Here we use optimal control theory to explain perceptual decision-making from a new perspective. Previous decision-making models, including drift diffusion, urgency-gating, recurrent neural networks, and dynamic field theory, typically fit parameters to match the data without considering how objectives of the nervous system would tune deliberation.^18,19,21,35,36^ As an example, for drift diffusion models the accumulation rate and decision threshold are often fit for each experimental condition.^19,21^ Conversely, we use cost function parameters that relate higher-level objectives of desirability and cognitive effort to deliberation. The control policy then tunes evidence accumulation and urgency in an emergent and principled manner based on biologically plausible objectives. It has been suggested that decision-making may consider an objective of reward rate,^2,37^ or relatedly a time cost.^27^ For example, Drugowitsch and colleagues have proposed a collapsing decision threshold for the drift-diffusion model that is a function of the time cost to accumulate evidence.^27^ Their proposed cost function modulates the decision threshold, but unlike our approach does not modulate how to tune deliberation itself. Others have used Bayesian models, but they require a mapping of each state to a probability and an associated cost to make an optimal selection.^38,39,40,41^ Conversely, we have a decision control policy where the decision emerges based on the inputted evidence and deliberation dynamics that are determined from biologically plausible objectives.

The optimal control theory we use, linear-quadratic regulator (LQR), allows for a principled framework to consider how biologically plausible objectives relate to behaviour.^13,14^ In this paper, we used an infinite-horizon solution to estimate the control gains to explain past work. Recent work has shown that perceptual decision-making is influenced by imposed decision deadlines.^28^ However a finite-horizon solution can be used to explain behavioural data with imposed time deadlines (**Supplementary A**). Other work has also examined differences in behaviour and neural activity when there is a free response time or forced viewing time,^25^ which our model is also able to replicate (**Supplementary B**). Throughout this paper we have modeled the influence of visual evidence on perceptual decision-making, however our model would readily be able to account for other sensory modalities^42^ or multisensory integration.^43,44^ Further, there is evidence to support the idea that the nervous system uses novel evidence (i.e., 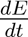) to make decisions,^2,45,46^ which could also be implemented with our modeling approach. Our computational framework proposes a cost function compatible with prominent theories of sensorimotor control.^13,14^ To replicate past work on embodiment,^46,47,48^ future work could create a composite computational framework where deliberation and motor control are governed by the same objectives.

Decision-making deficits may be seen in a different light with our approach. For example, those with Parkinson’s disease have deficits to the basal ganglia,^49^ which is implicated with reward-based processes. Using our framework, one can manipulate the desirability term (Q) to study the outcome of altered reward-based processing on decisions to mimic a neurological disorder. Deficits in cognitive processing, such as in those with mild cognitive impairment, aging, or other conditions that cause cognitive fatigue, could potentially relate to changes in the deliberation dynamics (A,B). Consequently, there are several exciting avenues to mechanistically study impaired decision-making and a host of neurological conditions.^50^

By considering biologically plausible objectives of cognitive effort and desirability, several decision-making behaviours simply emerge from our model. The model’s decision variable also shows a close resemblance to low-dimensional neural activity observed in non-human primates. Our new framework offers a theoretical bridge to link perceptual decision-making with neuroeconomics and sensorimotor neuroscience.

## Methods

Here we use the well-established, infinite-horizon linear quadratic regulator control frame work to simulate the decision-making process. First, we describe the dynamics and states for a single option (i). The deliberation dynamics are,

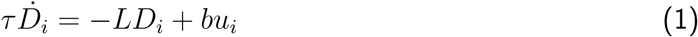

where *τ* is the time constant which governs the rate of change of the decision variable, *D* is the decision variable, *L* is the leak or forgetting term, *b* is the transformation of control signal to decision variable. We set both *b* and *τ* to 1 for all simulations. The neural control signal (**u***_i_*) drives the deliberation process as a function of the current states.

The states considered are,

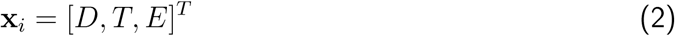

where *D* is the decision variable for the current option, *T* is the decision threshold, and *E* is the evidence for that specific option.

First, the dynamics were transformed into a system of first order differential equations and discretized. We then add state noise *σ_k_*.

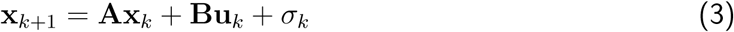

Here, *k* is the current time. The matrix **A** describes the deliberation dynamics and containing *L* and *τ*. **B** is a matrix that maps the control signal to the states and contains *b* and *τ*. The state noise has a mean of 0 and a covariance matrix with [*σ_D_, σ_T_, σ_E_*] on the diagonal and 0 elsewhere. The matrices are fully defined in **Supplementary C**.

We define an infinite horizon quadratic cost function with a state cost (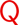), a control cost (R), and an interaction cost (S) between the states and the control signal.

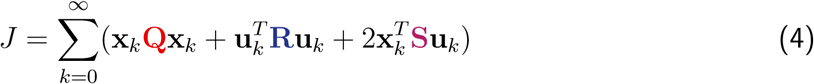

The cost function can be rewritten in matrix form as:

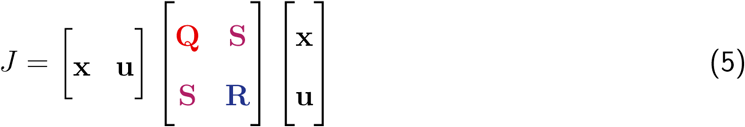

To expand to multiple potential options, we augment the state and control vectors as follows:

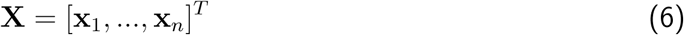

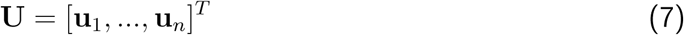

The term *n* represents the number of possible options. To simulate with cross-inhibition, we set a nonzero offdiagonal term in the 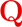 matrix for each decision variable corresponding to the other decision variables. The cost function matrix is fully described in Supplementary C.

Here, we approximate the optimal feedback policy as a function of the cost function and given dynamics using the algebraic Ricatti equation. The optimal control signal **U_k_** is equal to the control gains multiplied against the current states.

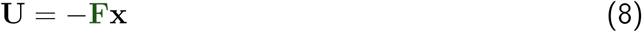

The control gain is found as a function of **P**,

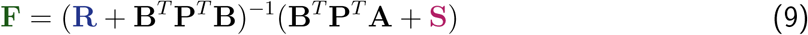

P is the solution to the discrete algebraic Ricatti equation as follows:

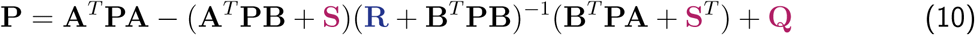

To approximate the infinite-horizon solution, we calculate the solution to the finite-horizon dynamic algebraic Ricatti equation for 100,000 time steps.

### Simulation

For each trial of a decision, we simulate all of the states until one of the decision variables crosses a threshold. Each trial is simulated with a time step of 1 ms. All parameters were selected to visually match experimental data and are shown in **Table 1**. Notably, differences between experimental manipulations (e.g., speed vs. accuracy instructions) are due to objective changes.

**Table 1:** Parameters for each simulation.

| Experiment | $L$ | $Q^D$ | $R$ | $Q^{Cross}$ | $\alpha$ | $\sigma^D$ | $\sigma^T$ | $\sigma^E$ |
| --- | --- | --- | --- | --- | --- | --- | --- | --- |
| Coherence | 3 | 0.013 | 0.00024 | 0.001 | 5.416667 | 0.02 | 0.0015 | 0.07 |
| Hick's Law | 3 | 0.0058 | 0.00024 | 0.003 | 41.66667 | 0.008 | 0.0015 | 0.01 |
| Coherence:<br>Accuracy Instructions | 3 | 0.08 | 0.00024 | 0.0011 | 6.666667 | 0.02 | 0.0015 | 0.07 |
| Coherence:<br>Speed Instructions | 3 | 0.012 | 0.00024 | 0.0011 | 6.666667 | 0.02 | 0.0015 | 0.07 |
| Tokens Task | 3 | 0.005 | 0.00024 | 0.002 | 4.583333 | 0.0045 | 0.0015 | 0.01 |
| Neural Activity | 3 | 0.015 | 0.00024 | 0.011 | 9.583333 | 0.03 | 0.0015 | 0.05 |
| Choking Under Pressure | 4 | **eq. 13** | 0.00024 | 0.0006 | 5.208333 | 0.0065 | 0.0015 | 0.07 |

#### Skewed Response Times and the Random Dot Motion Discrimination Task

To simulate the random dot motion discrimination task, we simulated 2 decision variables in parallel. The decision threshold (*T*) was constant at 1. We used the following coherences as the input evidence into the model [−0.5, −0.35, −0.25, −0.15, −0.1, −0.05, 0.05, 0.1, 0.15, 0.25, 0.35, 0.5]. *E*_1_ was set as the input evidence and *E*_2_ was equal to *−*1 *· E*_1_. For the skewed response time distribution (**Fig. 1C**), we displayed the response time distribution for the 0.15 coherence condition. The simulated behaviour for all of these coherences are shown in **Fig. 2A,B**. By changing only *Q_D_*, we can simulate the differences found between the accuracy (**Fig. 1B reddish purple, 2C**) and speed (**Fig. 1B purple, 2D**) instructions.

#### Hick’s Law

To simulate Hicks law, we modelled a decision variable for each possible option (**Fig. 1D**). The decision threshold was constant at 1. At time 0 ms, the evidence was set to 1 for the correct option.

#### Tokens Task

For time-varying evidence, we simulated 2 decision variables. We ran the simulation over 3000ms. The decision threshold (*T*) was constant at 1. The input evidence (*E*) was defined as

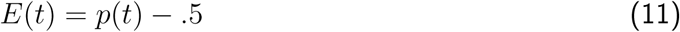

where *p*(*t*) is the probability at each timestep of an option. The probability (*p*(*L*)) of the left option being correct given number of tokens was defined as

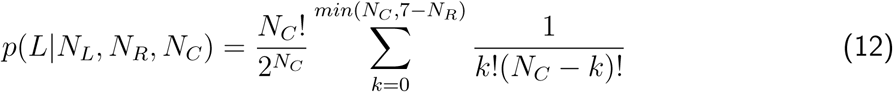

where *N_L_* is the number of tokens in the left target, *N_R_* is the number of tokens in the right target, *N_C_* is the number of tokens in the center. The probability of the right option being correct is 1 *− p*(*L*).

The tokens moved every 200 ms and there were 15 total tokens. For the ambiguous token patterns, the first 6 tokens alternated between the left and right to maintain a success probability close to 50%. The remaining tokens had an 80% probability towards the correct target.

In the bias-for token patterns the first 3 tokens moved towards the correct target, the next 3 tokens moved towards the incorrect target, and the remaining tokens moved with an 80% probability towards the correct target. In the bias-against token patterns, the first 3 tokens moved towards the incorrect target, the next 3 tokens moved towards the correct target, and the remaining tokens moved with an 80% probability towards the correct target. For the easy token patterns, each token had an 80% probability towards the correct target.

#### Neural Activity

For the neural activity in Steinemann (2024), we simulated 2 decision variables.^16^ We used the following possible coherences [−0.512, −0.256, −0.126, −0.064, -.032, 0.0, 0.032, 0.064, 0.126, 0.256, 0.512] as input evidence. *E*_1_ was set as the input evidence and *E*_2_ was equal to *−*1 *· E*_1_. The decision threshold was set to 1 beginning at 200 ms in the trial. To match the neural data, we set the initial state of the decision variable as 0.4.

#### Choking Under Pressure

For the choking-under-pressure phenomenon,^29^ we simulated 8 decision variables. The decision threshold was constant at 1. At 0 ms, we set the evidence for the correct option to 1.

Here we defined *Q^D^* as function relative to the average reward and current reward (*V_curr_*) of the trial. The rewards used in this experiment were 0.0, 0.1, 0.2, and 2.0 mL of juice with relative frequencies of 0.3167, 0.3167, 0.3167, and 0.05, respectively. Thus, the average reward (*V̅*) was 0.195. In line with prospect theory,^30^ the following equation is conceptually similar to a shift in reference point.

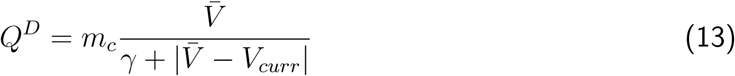

Here, the desirability for is a function of the average reward. Effectively, a different average reward would shift the reference of the relative desirability function. We set *γ* = 0.45 and *m_c_* = 0.075.

## Acknowledgements

We would like to thank Christopher Fetsch for his valuable comments on this manuscript.

