## Supplementary for "Deliberation is a controllable process governed by desirability and cognitive effort"

### Supplementary A: Finite Horizon

In the main text, we approximated an infinite horizon solution to the linear quadratic regulator controller. By using an infinite horizon solution to the optimal feedback gains, we found time-invariant control gains that are held constant to simulate decision-making behaviour. It is also possible to use the finite horizon linear quadratic regulator solution to simulate decision-making behaviour. Using the finite-horizon linear quadratic regulator solution, we replicate the results shown in **Fig 2A-B**. We capture the same decrease in response time (**Fig. S1A**) and increase in correct selection proportion (**Fig. S1B**) seen in the data and the infinite horizon solution.

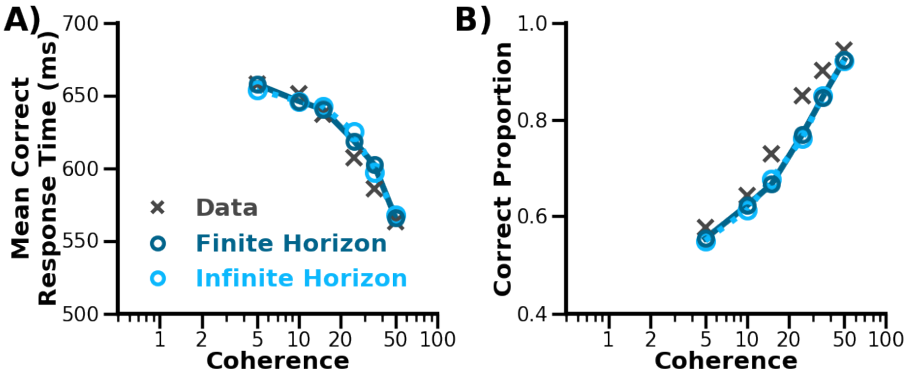


**Figure S1: Replication of Decision-Making Phenomena with Finite Horizon and Infinite Horizon solutions.** Participant behaviour (grey x's) from the random-dot motion discrimination task showing **(A)** the increase in correct responses and **(B)** the decrease in mean response time of correct responses with the stronger evidence (i.e. higher coherence) [Adapted from Ratcliff, 2008]. Both the infinite horizon solution (light blue; also shown in Fig. 2A) and finite horizon solution (dark blue) capture the behavioural trends.

To find the finite-horizon solution, we use a finite discrete linear-quadratic regulator. We define a standard quadratic cost function with a $(\mathbf{Q_N})$ terminal cost, running state cost $(\mathbf{Q})$, and control costs $(\mathbf{R})$. Note that $\mathbf{Q_k}$ was constant for all time steps. N is the total number of time steps in the trial.

$$J = \mathbf{x}_N^T \mathbf{Q}_N \mathbf{x}_N + \sum_{k=0}^{N-1} (\mathbf{x}_k \mathbf{Q}_k \mathbf{x}_k + \mathbf{u}_k^T \mathbf{R} \mathbf{u}_k + 2\mathbf{x}_k^T \mathbf{S} \mathbf{u}_k) \tag{1}$$

Here, we use the same parameters as the infinite-horizon solution (**See Table 1**) with $Q_N^D = 10$. We then used the discrete-algebraic Ricatti equation to solve for the time-varying feedback control gains $(F_k)$.

$$\mathbf{u}_k = -\mathbf{F}_k \mathbf{x}_k \tag{2}$$

The finite-horizon solution results in time-varying control gains (**Fig S2**). As expected, the time-varying control related to the decision variable (**Fig S2A**) and decision threshold (**Fig S2B**) gains rapidly increase closer to the time horizon. These specific parameters are related to minimizing the distance between the decision variable and decision threshold. For the finite-horizon, the deliberation is only permitted to continue for a set amount of time (e.g., imposed decision deadline during an experiment). Here we show that the finite-horizon and infinite-horizon control gains can be identical during the initial period. That is, one can consider the infinite-horizon solution to effectively be a finite-horizon solution with a long time scale. However, the finite-horizon control gains can change close to the imposed deadline, whereas the infinite horizon control gains are always held constant. For the observed behaviour shown in **Fig. S1**, the correct mean responses were within 700 ms. Thus, the two solutions resulted in identical

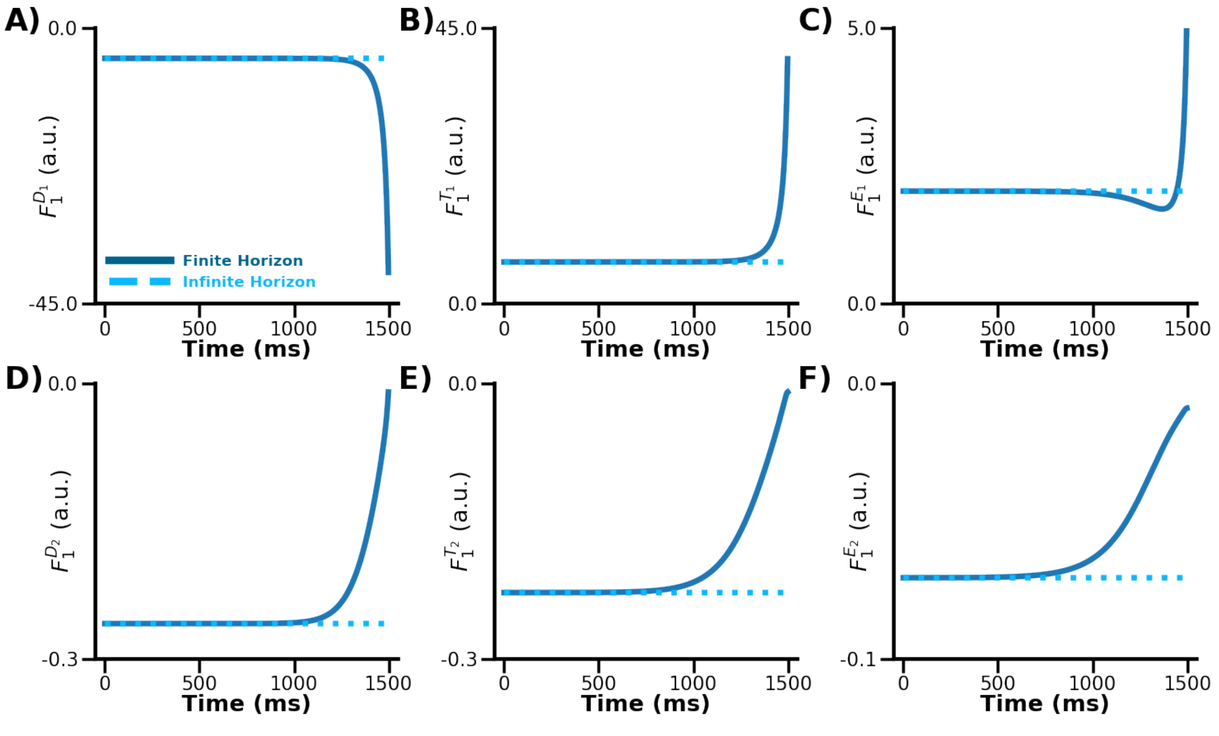


**Figure S2: The finite horizon and infinite horizon control gains over time.** The control gains that determine the control signal for option 1 based on **A)** option 1 decision variable, **B)** option 1 threshold, **C)** option 1 evidence, **D)** option 2 decision variable, **E)** option 2 threshold, and **F)** option 2 evidence. The control gains for the finite horizon (solid dark blue line) and the infinite horizon (dashed light blue line) formulations are shown to be similar here for the first 1000 ms. Here, we display the control gain multiplied by -1 to account for the negative shown in **equation 2**.

response times and decision times for this experiment, since the finite-horizon control gains are the same as the infinite-horizon prior to this time.

Importantly, using a finite-horizon solution we are able to replicate recent decision-making behaviour with an experimentally imposed decision deadline. Kira and colleagues (2025) modified the random-dot motion-discrimination task by implicitly imposing a decision deadline. Participants were asked to fixate their gaze on a random dot motion cloud while deciding on the movement direction of the dots. Participants made a saccade to the target that corresponded to movement of the dots. For phase 1, the participants were allowed to freely make decisions. For phase 2, researchers interleaved trials without a time deadline and trials with a randomly

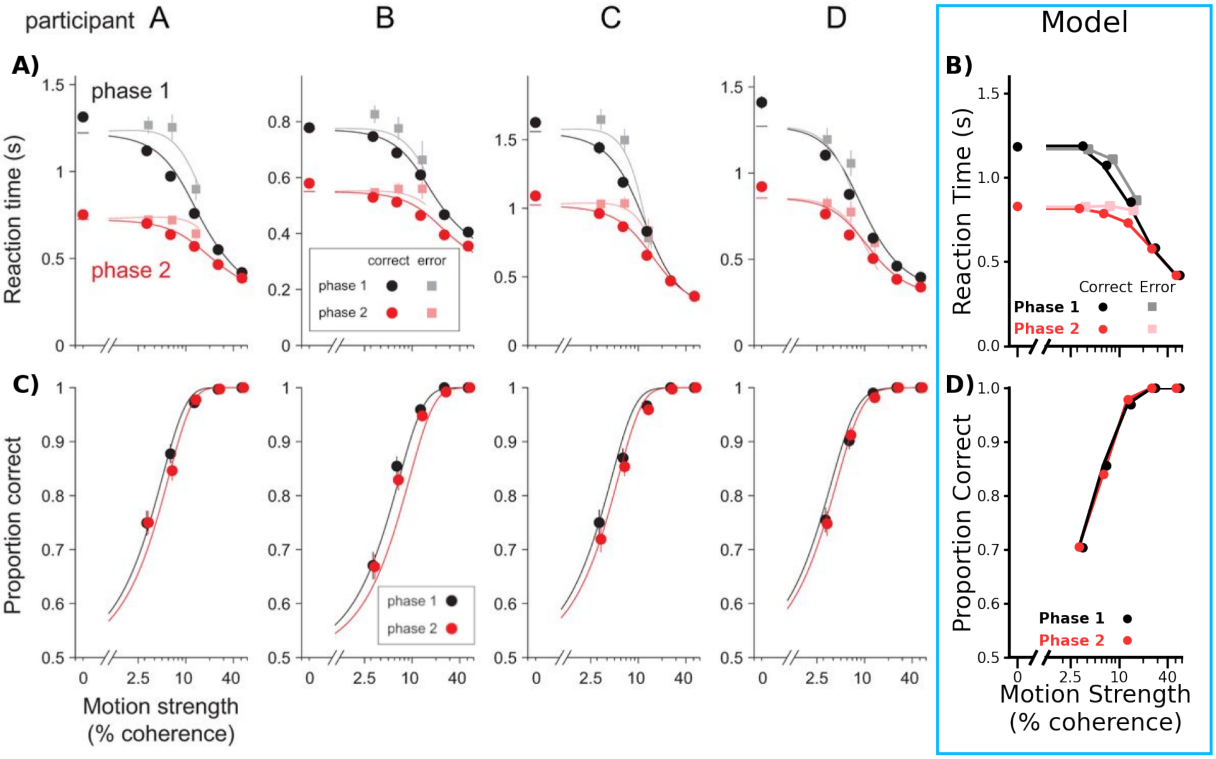


**Figure S3: Decision-making behaviour with and without a deadline.** Behaviour of four participants from the random-dot motion discrimination task without (phase 1) and with a time deadline (phase 2) [Adapted from Kira, 2025]. Across both phases, the participants made **A)** faster and **C)** more correct decisions with stronger coherences. **A)** Mean reaction time for the correct (circles) and error (squares) responses for each coherence. Participants had earlier correct (red circle) and error (pink square) reaction times in phase 2 compared to the correct (black circle) and error (grey square) reaction times in phase 1. **C)** Participant proportion of correct responses for each coherence in phase 1 (black) and phase 2 (red). We used a long finite-horizon for phase 1 (black circle and grey square) and a short finite-horizon solution for phase 2 (red circle and pink square) to simulate reaction times and correct proportions given different deadlines. **B)** Here, our model replicates the earlier decision times found in phase 2 compared to phase 1. **D)** Similar to the behaviour, our model also find more correct responses with higher coherence.

selected time deadline. Researchers compared the reaction times (**Fig. S3A**) and the correct response proportions (**Fig. S3C**) between the trials in phase 1 and the trials in phase 2 without a time deadline. They found earlier decisions in phase 2 compared to phase 1. Here, we simulate phase 1 using a finite-horizon solution with a very long decision deadline (i.e., 10,000 ms) and phase 2 using a much shorter decision deadline (i.e. 1,500 ms). Our model captures the similar trends for the reaction times (**Fig. S3B**) and the correct response proportions (**Fig. S3D**) as a function of coherence. In alignment with behaviour, our model finds earlier decisions in phase 2 (**Fig. S3AB**, red) compared to phase 1 (**Fig. S3AB**, black).

### Supplementary B: Fixed Viewing Duration and Free Response

Prior work has measured the neural activity of non-human primates during a random dot-motion discrimination task during a free response and a fixed viewing duration condition. In the free response condition, the non-human primate made a saccade to a target as soon as they made

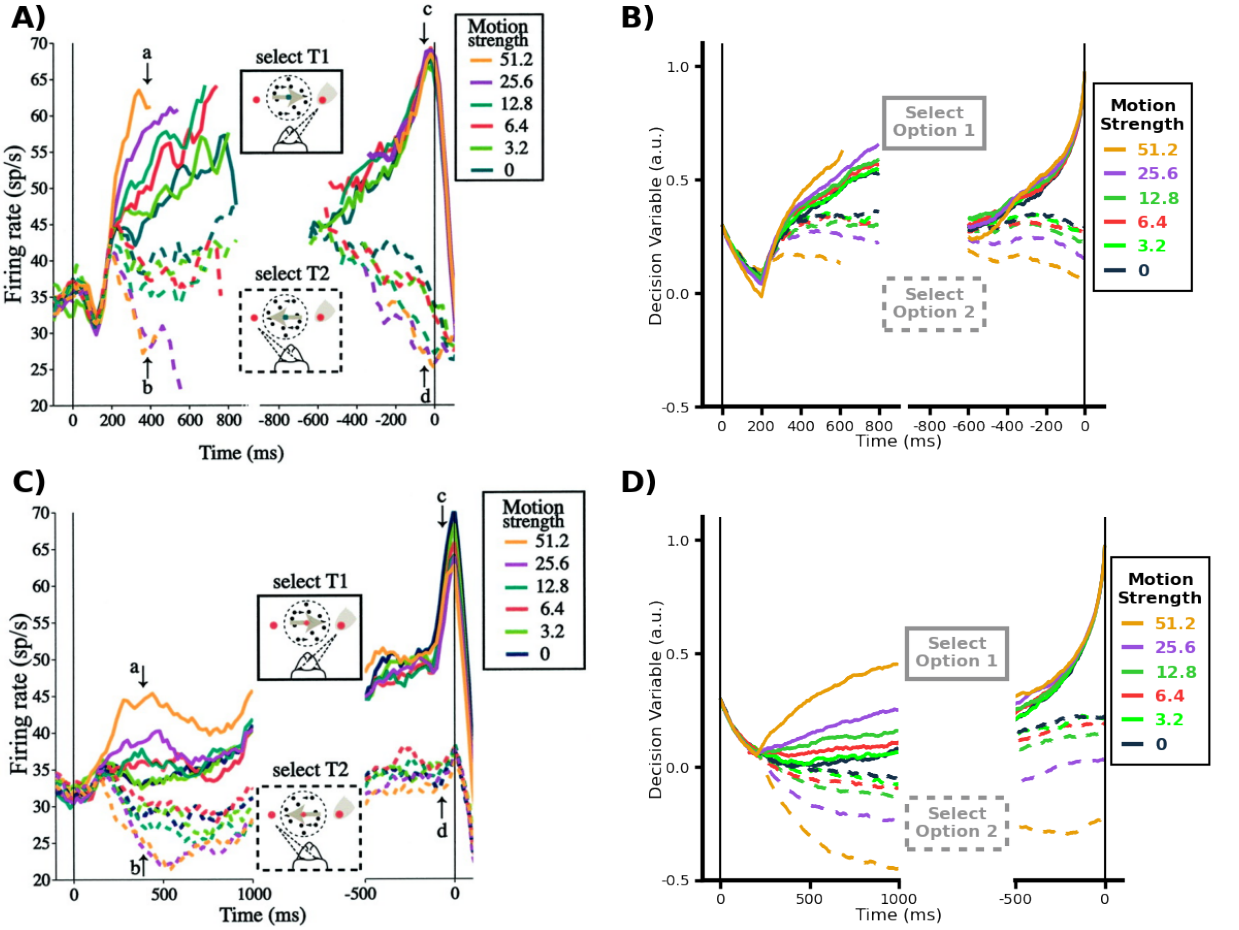


**Figure S4: Neural Activity of during Reaction Time and Fixed Duration perceptual decision-making tasks.** **A,C)** Neural Activity in the lateral intraparietal sulcus of a non-human primate during a random dot motion discrimination task **A)** in a reaction time condition and **C)** fixed duration condition [Adapted from Roitman (2002)]. **A)** In the reaction time condition, the monkey was free to respond as soon as they made a decision. **C)** In the fixed duration condition, the monkey was forced to view the evidence for 1000 ms. **B,D)** Simulated model dynamics for a reaction time and fixed duration perceptual decision-making task. **B)** Average simulated decision variable over time of one potential option in the reaction time condition. **D)** Average simulated decision variable over time of one potential option in the fixed duration condition. The decision variable does well to capture the trends of temporally evolving neural activity across both conditions.

their decision. Aligned with the instructions, the neural activity has an evidence-independent increase (**Fig. S3A**) across the coherences. In the fixed viewing duration condition, they were forced to view the random dot-motion for at 1000 ms, the stimulus would disappear, and they would then make a saccade. Unlike the free response condition, there is no evidence-independent increase in neural activity across the coherences (**Fig. S3C)**.

Here, we are able to model both the free response and fixed duration conditions using our computational framework. To simulate the free response condition, we used a similar paradigm set up as **Neural Activity** in the **Methods**. For the free response condition, we set the decision threshold to 1 at 200ms. Here, we replicate the increase shown in the behaviour due to urgency (**Fig. S4B**). The only change required to simulate the fixed duration condition was to set the decision threshold to 1 at 1000 ms. Because the decision threshold is set to 0, there is no influence of urgency. In line with the neural data, the evolution of the decision variable is primarily due to evidence and is centered around 0 (**Fig. S4D**). Here, we capture the differences between the free response and fixed duration conditions due to the emergent properties of urgency in our model.

#### Supplementary C: Full Model and Model Sensitivity Analysis

Here, we expand on the dynamic equation for a deliberation process with two options. As described in the methods, this can be easily expanded to any number of options.

$$\mathbf{X}_{k+1} = \mathbf{A}\mathbf{x}_k + \mathbf{B}\mathbf{U}_k + \boldsymbol{\epsilon}_k \tag{3}$$

where the $X$ is a vector of the states, $\mathbf{U}$ is the vector of the control signals, $\mathbf{A}$ is a matrix that describes the dynamics, $\mathbf{B}$ describes the transformation of control signals to states, and $\boldsymbol{\epsilon}$ is a vector of the noise for each state.

$$\mathbf{A} = \begin{bmatrix} 1-\frac{L}{\tau}h & 0 & 0 & 0 & 0 & 0 \\ 0 & 1 & 0 & 0 & 0 & 0 \\ 0 & 0 & 1 & 0 & 0 & 0 \\ 0 & 0 & 0 & 1-\frac{L}{\tau}h & 0 & 0 \\ 0 & 0 & 0 & 0 & 1 & 0 \\ 0 & 0 & 0 & 0 & 0 & 1 \end{bmatrix}, \mathbf{B} = \begin{bmatrix} \frac{b}{\tau}h & 0 \\ 0 & 0 \\ 0 & 0 \\ 0 & \frac{b}{\tau}h \\ 0 & 0 \\ 0 & 0 \end{bmatrix}, \mathbf{X}_k = \begin{bmatrix} D_k^1 \\ T_k^1 \\ E_k^1 \\ D_k^2 \\ T_k^2 \\ E_k^2 \end{bmatrix} \mathbf{U}_k = \begin{bmatrix} u_k^1 \\ u_k^2 \end{bmatrix} \boldsymbol{\epsilon} = \begin{bmatrix} \epsilon_k^{D^1} \\ \epsilon_k^{T^1} \\ \epsilon_k^{E^1} \\ \epsilon_k^{D^2} \\ \epsilon_k^{T^2} \\ \epsilon_k^{E^2} \end{bmatrix}$$

We also expand on the cost function for two options.

$$J = \left[\begin{array}{c|c} \mathbf{X}^T & \mathbf{U}^T \end{array}\right] \left[\begin{array}{c|c} \mathbf{Q} & \mathbf{S} \\ \hline \mathbf{S} & \mathbf{R} \end{array}\right] \left[\begin{array}{c} \mathbf{X} \\ \hline \mathbf{U} \end{array}\right] \tag{4}$$

$$J_k = \left[\begin{array}{cccccc|cc} D_k^1 & T_k^1 & E_k^1 & D_k^2 & T_k^2 & E_k^2 & u_k^1 & u_k^2 \end{array}\right] \left[\begin{array}{cccccc|cc} Q^D & -Q^D & 0 & Q^{Cross} & 0 & 0 & 0 & 0 \\ -Q^D & Q^D & 0 & 0 & 0 & 0 & 0 & 0 \\ 0 & 0 & 0 & 0 & 0 & 0 & -S & 0 \\ Q^{Cross} & 0 & 0 & Q^D & -Q^D & 0 & 0 & 0 \\ 0 & 0 & 0 & -Q^D & Q^D & 0 & 0 & 0 \\ 0 & 0 & 0 & 0 & 0 & 0 & 0 & -S \\ \hline 0 & 0 & -S & 0 & 0 & 0 & R & 0 \\ 0 & 0 & 0 & 0 & 0 & -S & 0 & R \end{array}\right] \left[\begin{array}{c} D_k^1 \\ T_k^1 \\ E_k^1 \\ D_k^2 \\ T_k^2 \\ E_k^2 \\ \hline u_k^1 \\ u_k^2 \end{array}\right] \tag{5}$$

Here, the red relates to the objective of desirability, blue relates to the objective of cognitive effort, and purple relates to the cost of parallel decision variables. After performing the matrix multiplication, we end up with the equation:

$$J_k = \sum_{i=1}^{n} \left[ Q^D (D_k^i)^2 + Q^D (T_k^i)^2 - 2Q^D (D_k^i)(T_k^i) + R(u_k^i)^2 + -2S(E_k^i)(u_k^i) \sum_{\substack{j=1 \\ j \neq i}}^{n} + Q^{Cross}(D_k^i)(D_k^j) \right] \tag{6}$$

Here, we define $S = \alpha R$. The S is the interaction cost between the neural control signal and evidence. Thus, we suggest that the interaction cost is a function of the weighting on cognitive effort.

$$J_k = \sum_{i=1}^{n} \left[ (Q^D {D_k^i}^2 - 2Q^D D_k^i T_k^i + Q^D {T_k^i}^2) + (R{u_k^i}^2 - 2\alpha R E_k^i u_k^i) + \sum_{\substack{j=1 \\ j \neq i}}^{n} Q^{Cross}(D_k^i)(D_k^j) \right] \tag{7}$$

$$J_k = \sum_{i=1}^{n} \left[ Q^D ({D_k^i}^2 - 2D_k^i T_k^i + {T_k^i}^2) + R({u_k^i}^2 - 2\alpha E_k^i u_k^i) + \sum_{\substack{j=1 \\ j \neq i}}^{n} Q^{Cross}(D_k^i)(D_k^j) \right] \tag{8}$$

$$J = \sum_{i=1}^{n} [ Q^D (D_k^i - T_k^i)^2 + R(u_k^i - 2\alpha E_k^i u_k^i) + \sum_{\substack{j=1 \\ j \neq i}}^{n} Q^{cross} D_k^i D_k^j ] \tag{9}$$

As a reminder, the red relates to the objective of desirability, blue relates to the objective of cognitive effort, and purple relates to the cost of parallel decision variables. These components of the cost functions lead to urgency, evidence accumulation, and cross-inhibition.

We have performed a model sensitivity analysis. From the standard random-dot motion discrimination task, we simulated 0.15 coherence. We used the parameters that replicated the behaviour in **Fig. 2A-B** as the baseline. We varied each parameter individually, while holding every other parameter constant. Corresponding mean correct response time and selection proportion are shown in **Fig. S5**.

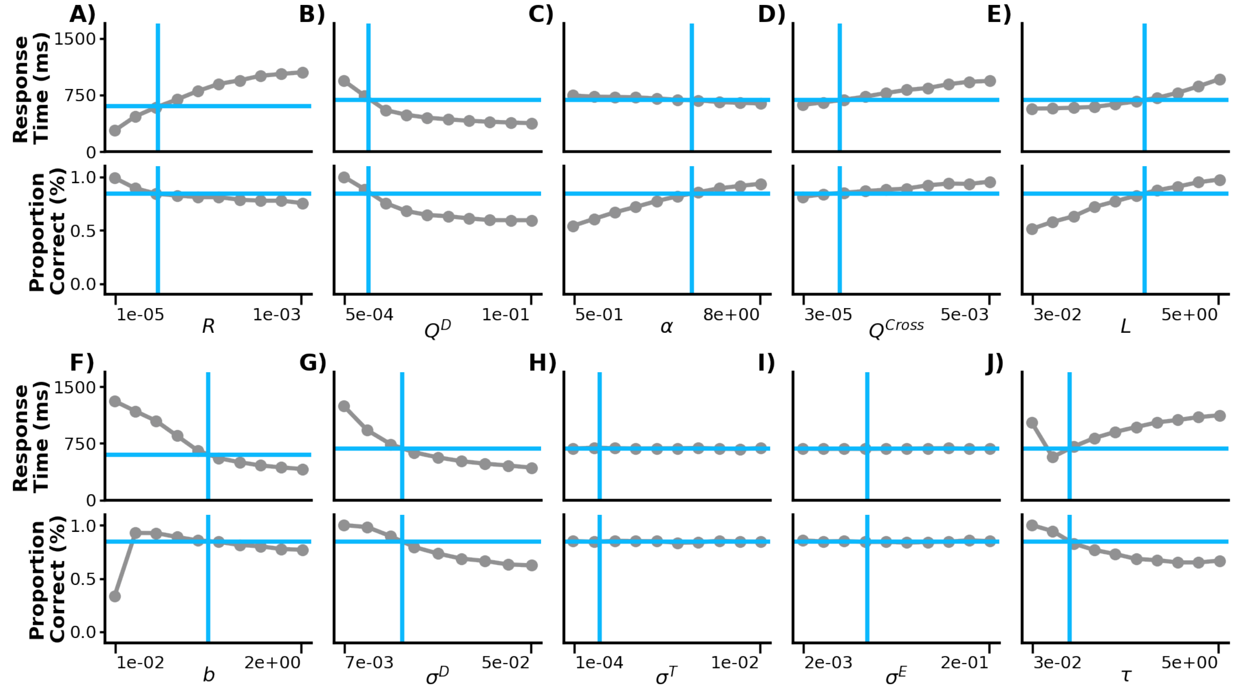


**Figure S5: Sensitivity Analysis.** We performed a one-factor-at-a-time sensitivity analysis. That is, we changed one parameter while holding the other parameters constant. We tested **A)** $R$, **B)** $Q^D$, **C)** $\alpha$, **D)** $Q^{Cross}$, **E)** $L$, **F)** $b$, **G)** $\sigma^D$, **H)** $\sigma^T$, **I)** $\sigma^E$, **J)** $\tau$ For each free change in parameter, we found the response time and proportion of correct selections in the 0.15 coherence condition shown in **Fig 2A,B**. The vertical blue line is the model parameter used in this main study. The horizontal blue line is the outcome measure from the selected parameters.
